# Who Funds Open Data Sharing? Analysis of data availability statements in biomedical publications

**DOI:** 10.64898/2026.07.17.739022

**Authors:** Josh Lawrimore, Christoph Li, Dustin Moraczewski, Jean-Baptiste Poline, Adam Thomas

## Abstract

Open data sharing is increasingly mandated by research funders, journals, and institutions, yet large-scale compliance measurement remains challenging. We analyzed 951,949 open access biomedical research articles published between January 2024 and June 2025 using a dual-source pipeline: PDF-based text extraction (MinerU) where PDFs were available and PMC XML otherwise, followed by algorithmic detection of data sharing statements (oddpub v7.2.3), enriched with funder, journal, and institutional metadata from OpenAlex. The corpus included 294,172 PDF-covered articles (30.9%) and 657,777 XML-only articles (69.1%). We found an overall open data rate of 8.7%, rising to 11.7% among funder-linked articles (those with at least one funder identified in the metadata). Rates varied more than tenfold across the research ecosystem: leading major funders reached observed open data rates of 20–24%, while top journals reached observed rates of 70–86%, with corrected estimates as high as 92.9% (Nature Genetics) after adjusting for XML-only coverage limitations. PDF-based detection identified approximately 52% more data sharing statements than XML-based methods on the same articles. These observed rates differ markedly across funders and journals, and current overall sharing remains far below universal compliance. These patterns provide an empirical baseline against which future policy changes can be measured. An interactive dashboard at https://www.opensciencemetrics.org enables stakeholders to explore and benchmark these results.

## 1 Introduction

The open science movement has placed data sharing at the center of efforts to improve the transparency and reproducibility of research. The FAIR principles, that research data should be Findable, Accessible, Interoperable, and Reusable, have become a widely endorsed framework for scientific data stewardship (*1*). Meanwhile, surveys consistently reveal that a majority of researchers have encountered difficulties reproducing published results, underscoring the practical importance of access to underlying data (*2*). Studies have also demonstrated that publications linking to shared datasets receive more citations, creating incentive structures that complement institutional mandates (*3*).

In response, data sharing has become a shared priority of biomedical research funders worldwide, anchored by policies from the National Institutes of Health (*4*), the European Commission (*5*), UK Research and Innovation (*6*), the Wellcome Trust (*7*), the Deutsche Forschungsgemeinschaft (*8*), the Swiss National Science Foundation (*9*), the Agence Nationale de la Recherche (*10*), and the US National Science Foundation (*11*), as well as multilateral frameworks including the FAIR Data Principles (*1*) and the UNESCO Recommendation on Open Science (*12*). This policy convergence has not been matched by measured practice. In the most comprehensive assessment to date— 2.75 million open access PubMed Central articles published through 2020—Serghiou et al. found that the prevalence of data sharing, while rising over time, had reached only an estimated 15% of all articles, and 17% of research articles, by 2020, despite the policies already in force by then (*13*). Funders and journals differ substantially in the stringency, scope, and enforcement of their data sharing policies, yet population-scale measurements of the open data rates that co-occur with these differences remain limited.

The scope and pace of these policies are generating new measurement demands. NIH’s $1M Data Sharing Index (“S-index”) Challenge, whose Phase 1 finalists were named in September 2025 (*14*, *15*), seeks a researcher-level metric analogous to the h-index for data sharing; operationalizing such a metric at population scale will require exactly the kind of corpus-wide measurement infrastructure we describe here.

Journals have also emerged as influential gatekeepers of data sharing. Many high-impact journals now require data availability statements or deposit of data in public repositories as a condition of publication. Nature Research journals, Cell Press titles, and PLOS journals have progressively strengthened their data sharing policies over the past decade. However, the relationship between policy existence and actual compliance is not straightforward: previous work has shown that data availability statements do not always correspond to genuinely accessible data (*16*).

Measuring actual data sharing at scale is essential for tracking compliance with these policies and for benchmarking funders, journals, and disciplines against one another. Without systematic measurement, funders cannot benchmark their grantees’ compliance, journals cannot assess the impact of editorial policies, and institutions lack the data needed to support their researchers. Yet scalable measurement presents significant methodological challenges. Manual assessment of data sharing is labor-intensive and impractical for large corpora (*17*). Text-mining approaches, while scalable, require access to the full text of the publication. The XML versions of articles in the PubMed Central Open Access (PMCOA) collection may miss data sharing statements present in final published versions: data sharing info is often added late in the publication process or appears in sections omitted from the XML.

In this work we address these gaps by using a PDF-first detection pipeline. Full-text PDFs are processed using MinerU (*18*), a machine learning-based document extraction tool, and the resulting text is analyzed by oddpub v7.2.3 (*19*), an algorithm that identifies open data and open code sharing statements through pattern matching. This combination captures data sharing statements that XML-based methods miss, including those in supplementary sections, figure captions, and formatted data availability statements. On articles with both PDF and XML coverage, our PDF-based pipeline detects approximately 52% more open data statements than XML alone.

We analyzed 951,949 open access biomedical research articles published between January 2024 and June 2025, characterizing data sharing rates across 1,833 funders and 1,388 journals. Of these, 294,172 articles had PDF-derived full text and 657,777 were analyzed from XML only; journal-level correction factors from head-to-head PDF versus XML comparisons were applied to account for this differential coverage. We report observed open data rates (exact corpus proportions) alongside corrected estimates with 95% imputation intervals reflecting the uncertainty of the coverage correction. Our results reveal more than tenfold variation in data sharing rates across the research ecosystem, with clear signals that funder and journal policies are associated with substantially higher compliance.

All results are available through an interactive dashboard at https://www.opensciencemetrics. org, enabling funders, journal editors, institutional administrators, and researchers to explore the data across multiple dimensions.

## 2 Methods

### 2.1 Data Collection

We analyzed open access biomedical research articles with publication dates from 1 January 2024 through 25 June 2025. The corpus is anchored on the PubMed Central Open Access (PMC-OA) subset, which supplies all XML full text and 926,299 of the 951,949 articles (97.3%); the remaining 25,650 articles (2.7%) are open access records that are not deposited in PMC and enter the corpus through a publisher-hosted PDF alone. The closing date is not an analytic choice: it is the coverage end date of the 2025 mid-year PMC-OA baseline bulk package, the archival snapshot that defines the corpus and supplies its XML. Article-level metadata (including publication type, journal of publication, funder information, and author institutional affiliations) was obtained from OpenAlex (*20*), an open bibliographic database that aggregates records from Crossref, PubMed, institutional repositories, and other sources, retrieved via its public API on 17 December 2025 (OpenAlex is continuously updated and does not issue discrete versions; the metadata reflect its holdings on that date). We classified each article by type using the OpenAlex type field and the PMC XML `article-type`: research articles (OpenAlex `type = article` or PMC `article-type = research-article`, with OpenAlex taking precedence) were retained, and non-research records (reviews, editorials, letters, corrections, and errata) were excluded. This yielded a corpus of 951,949 research articles, each represented once and keyed by its PubMed identifier (PMID); the PMC-OA subset does not index preprints, so no deduplication of preprint–published-version pairs was required. Corpus-wide rates are computed over all 951,949 articles. The entity-level analyses necessarily cover subsets of this corpus: the journal analysis is restricted to the 778,534 articles appearing in the 1,388 journals that meet the minimum analysis threshold of 100 articles (Section 2.5.2), and the funder analysis to the 573,510 articles carrying at least one funder link in OpenAlex. Full-text PDFs were retrieved, where an open access copy could be located, from the open access locations recorded by OpenAlex—predominantly publisher-hosted versions of record, with a smaller contribution from author manuscripts, and only a negligible share from PMC itself (294,172 articles, 30.9%). The remaining 657,777 articles (69.1%) were analyzed using PMC XML full text. This distinction matters for interpretation: the PDF and XML arms differ not only in extraction method but in the version of the article they represent, since the publisher PDF is typically the version of record while PMC XML may omit late-added material.

### 2.2 Study Design

This is a cross-sectional, descriptive analysis. We compare observed open data rates across funders and journals within a single publication window using all open access PMC articles meeting our inclusion criteria. Because funders and journals are not randomly assigned to articles, and because funder, journal, and research discipline are tightly confounded in practice, the rate differences we report describe associations between entity-level context and open data rates; they do not identify the causal effects of any specific policy, mandate, or intervention. We make this scope choice deliberately: rigorous causal inference would require either a longitudinal pre/post design around a specific policy change or a quasi-experimental comparison with explicit controls, both of which we leave to future work. Statements throughout this paper about policy effectiveness, mandates “working,” or community norms “driving” open data behavior should be read as descriptive co-occurrence, not causal estimation.

### 2.3 PDF Processing and Open Data Detection

PDFs were processed using MinerU (*18*), a machine learning-based extraction tool optimized for scientific documents that preserves document structure including tables, figures, and supplementary sections. The extracted text was then analyzed using oddpub v7.2.3 (*19*), an algorithm that detects open data and open code sharing statements in scientific publications through pattern matching against a curated dictionary of data sharing phrases, repository names, and accession number formats. Each article was classified as containing (or not containing) an open data statement and an open code statement.

For articles without available PDFs, we used the PMC XML full text as the input to oddpub. This dual-source approach maximized corpus coverage while enabling head-to-head comparison of detection rates between the two extraction methods.

### 2.4 Metadata Enrichment

Funder names reported in OpenAlex were normalized using a curated alias mapping that aggregates related entities. For example, individual NIH institutes (e.g., NIMH, NIAID, NCI) were aggregated under their parent organization where appropriate, with child institute article counts deduplicated to avoid double-counting. Likewise, European Union research-funding programmes—Horizon 2020, Horizon Europe, the European Research Council, the Marie Skłodowska-Curie Actions, and the earlier Framework Programmes—were consolidated under the European Commission, since these are instruments of a single funder. This mapping was curated by the authors and is released with the analysis code. OpenAlex additionally surfaces a number of sub-agency programmes as separate funder entities whose parent agency also appears in the corpus (for example, NSF directorates and divisions, the DOE Office of Science and its national laboratories, and USDA’s National Institute of Food and Agriculture). Because neither the OpenAlex funder records nor our registry populate a parent-funder field for these entities, we resolved them against the Research Organization Registry (ROR) hierarchy: within a vetted set of root agencies, every corpus funder that ROR places beneath an agency at any depth was folded into that agency, with article counts deduplicated so that an article crediting both a programme and its agency is counted once. Degree-granting institutions were not folded into funders. Root agencies were restricted to those whose own sub-units OpenAlex surfaces separately, since ROR encodes administrative containment rather than the funder of record; for biomedical research the funder of record is the operating division (e.g., NIH, CDC) rather than the parent department, so these are reported as top-level funders. OpenAlex works counts were used to filter for established funders, excluding small or niche entities with fewer than 100,000 aggregated works from the main figures while retaining them in supplementary rankings. Funder–article links were refreshed from OpenAlex’s expanded 2026 funding data (retrieved 12 July 2026), which incorporates direct ingestion of funder grant records and full-text mining of acknowledgement sections in addition to Crossref-deposited metadata; the corpus and all per-article open data outcomes are unchanged from the December 2025 snapshot, so only the funder attribution reflects the later date.

Journal names were taken directly from the OpenAlex primary_location.source.display_name field. A small number of records carry an aggregator or index name rather than a genuine journal in this field; we excluded such non-journal sources (e.g. a “PubMed” pseudo-source) from the journal-level analysis, leaving 1,388 journals. These records remain in the corpus-level totals.

### 2.5 Quantitative Analysis

Open data rates were calculated as the proportion of articles with detected sharing statements, expressed as percentages. We use two corpus-wide reference rates. The *corpus-wide rate* (8.7%) is the open data rate across all articles in the corpus (all PMC-OA research articles in the analysis window) and is the reference for the journal analysis. The *funder-linked rate* (11.7%) is the rate across the subset of articles with at least one funder identified in the OpenAlex metadata, counting each article once regardless of how many funders it lists; it is the reference line in the funder figure. Because a given article may credit several funders, per-funder rates count that article in each of its funders’ totals, but the funder-linked reference counts it once. A missing funder link does not imply that the work was unfunded (see Results).

#### 2.5.1 Correction Factors for XML-Only Articles

Because PDF-based text extraction detects more data sharing statements than XML-based extraction, articles with only XML coverage have systematically underestimated open data rates. We developed a correction procedure using the subset of articles with both PDF and XML text (the “head-to-head” subset). For each journal with sufficient head-to-head data, we computed the “best” detection rate as the proportion of articles where *either* the PDF or XML extraction identified an open data statement. This best rate represents our estimate of the true open data rate for that journal, since it combines the complementary strengths of both extraction methods.

For articles with only XML coverage, we applied the journal-specific best detection rate as a correction factor. The corrected open data count for a given entity (funder or journal) was calculated

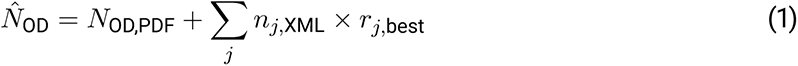

where *N*_OD,PDF_ is the observed open data count among PDF-covered articles, *n_j,_*_XML_ is the number of XML-only articles in journal *j*, and *r_j,_*_best_ is the best detection rate for journal *j* from the head-to-head subset. For journals without sufficient head-to-head data, a global fallback rate—the corpus-wide head-to-head best detection rate, 16.0%—was used in place of a journal-specific factor. This fallback is load-bearing: it applies to 738 of the 1,388 journals analyzed, which together hold 238,964 articles, or 31% of the journal-level corpus (778,534 articles). For these journals the correction is not informed by any within-journal head-to-head evidence, and the resulting corrected estimates should be read as the corpus-average expectation rather than as a journal-specific finding. Their imputation intervals reflect only the precision of the global rate, which is estimated over a large sample and is therefore narrow; that narrowness is a property of the global estimate and should not be read as confidence about any individual journal. The corrected estimate was floored at the observed count to ensure that corrections never reduce an already-observed signal.

Because this analysis is a census of all articles meeting our inclusion criteria rather than a sample drawn from a larger population, observed open data rates are exact proportions for the corpus and are reported without confidence intervals. The only quantity carrying statistical uncertainty is the XML-only correction, which *imputes* the open data statements that PDF-based processing would have detected in articles for which only XML was available. We quantify this uncertainty with a 95% *imputation interval*: a Wilson score interval (*21*) on the head-to-head best detection rate, propagated through the weighted correction of Equation (1). This interval should be read as the uncertainty introduced by the coverage correction, not as sampling uncertainty of the headline rate. Where the correction does nothing—an entity with no XML-only articles, or one whose imputed estimate is floored at the observed count—no imputation interval is reported.

#### 2.5.2 Figure Inclusion Thresholds

The number of articles per entity is heavily skewed, and the long tail of that distribution is populated largely by entities that are not comparable to the large, established organizations the figures are meant to characterize: narrowly scoped or one-off programmes, idiosyncratic grant labels, and artifacts of metadata parsing. Including them alongside major world funders would compare unlike things. To restrict each figure to entities of broadly comparable scope, we therefore applied Weibull-derived survival thresholds to entity-level article counts. A threshold is the article count at which the survival function of a Weibull distribution fitted to those counts equals a stated percentage; a lower survival percentage yields a higher article-count cutoff. For the main figures we used the 3% survival quantile for funders (at least 2,694 funder-linked articles) and the 2% survival quantile for journals (at least 4,262 articles); funders additionally required 100,000 aggregated OpenAlex works to exclude small or niche entities. The supplementary tables use a more permissive 5% survival quantile—at least 1,708 funder-linked articles (with a 50,000-works minimum) for funders, and at least 1,815 articles for journals. All entities above the minimum analysis threshold of 100 articles appear in the full supplementary rankings.

## 3 Results

### 3.1 Overall Open Data Rates

Across our corpus of 951,949 open access biomedical research articles published from January 2024 through June 2025, we detected open data sharing statements in 8.7% of publications. Among *funder-linked* articles—those for which OpenAlex identifies at least one funder—the rate was 11.7%. We caution that a missing funder link does not imply that the work was unfunded: such research is typically supported by institutional, self-, or other funding that is simply not captured in the available metadata, so this contrast reflects the presence of *identifiable* funding, not funding *per se*. Data sharing also increased with the number of distinct funders credited on an article, consistent with the collaborative and often mandate-bound nature of heavily co-funded research.

### 3.2 Variation by Funder

Open data rates varied substantially across the 1,833 funders analyzed, and which funders lead depends on how the set is filtered. We therefore distinguish two strata. Among the *major world funders* that account for the bulk of global research output—those exceeding both the Weibull article-count threshold and 100,000 aggregated OpenAlex works, shown in Figure 1—the leaders reached roughly twice the 11.7% funder-linked rate. Among *all* 1,833 funders meeting the minimum analysis threshold—most of which fall below the thresholds applied to Table 1 and appear only in the complete rankings released with the analysis code—the highest per-article rates were instead achieved by smaller, specialized research organizations, reaching up to four times that rate. Funder size and per-article open data rate are thus largely decoupled—a point we return to in the Discussion—so we describe each stratum in turn.

**Figure 1:**
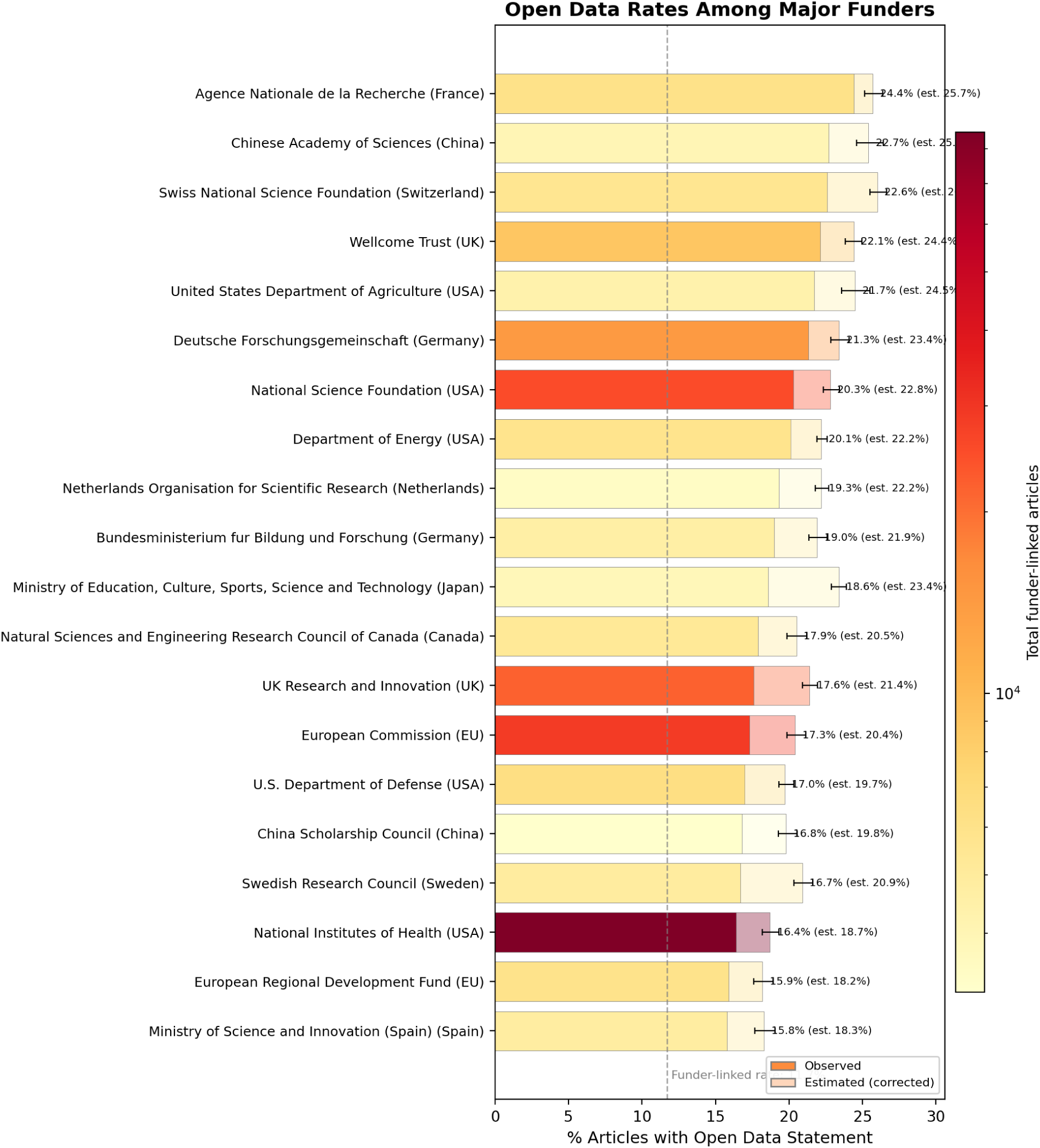
Open data rates among major biomedical research funders (2024–2025 research articles). Bar length shows the observed open data rate (full opacity) and the estimated corrected rate (lighter shade) after applying journal-level PDF vs. XML correction factors. Error bars indicate the 95% imputation interval on the corrected estimate (uncertainty in the XML-only coverage correction, not sampling uncertainty of the observed rate); entities where the correction does nothing carry no interval. Bar color encodes total funder-linked article count on a log scale (yellow = fewer, red = more). Dashed line shows the open data rate across all funder-linked articles (11.7%). The 20 highest-rate funders among those exceeding both the Weibull-derived 3% survival article-count threshold (≥2,694 articles) and 100,000 aggregated OpenAlex works are shown; rankings for the remaining funders in this stratum appear in Table 1, and the complete rankings for all 1,833 funders are released with the analysis code.

Among the major world funders shown in Figure 1, the Agence Nationale de la Recherche (France) led at 24.4% observed, followed by a cluster of large national funders near 22–23%: the Chinese Academy of Sciences (22.7%), the Swiss National Science Foundation (22.6%), and the Wellcome Trust (22.1%), with the U.S. Department of Agriculture (21.7%), the Deutsche Forschungsgemeinschaft (21.3%), and the U.S. National Science Foundation (20.3%) close behind. The U.S. National Institutes of Health—by far the largest funder in the corpus (85,144 funder-linked articles)—reached 16.4%, with UK Research and Innovation at 17.6% and the European Commission at 17.3%. Figure 1 shows the 20 highest-rate members of this stratum, all of which exceeded the 11.7% funder-linked rate. The stratum itself contains 35 funders, and six fall below that line—among them the Japan Society for the Promotion of Science (11.4% across 20,612 funder-linked articles), the seventh-largest funder in the stratum. Per-article rates thus vary widely even among major funders, and the largest funders are not the highest per-article performers.

Across the full set of 1,833 funders, the highest observed rates were concentrated among smaller, mission-focused research organizations. These fall below the article-count and works thresholds applied to Table 1 and Figure 1, which are restricted to major world funders; the entities named below therefore appear not in those exhibits but in the complete rankings released with the analysis code (Data Availability). The European Molecular Biology Laboratory (EMBL) recorded among the highest observed rates at 46.6% (104 of 223 articles), followed by the European Molecular Biology Organization (EMBO) at 41.0% (153 of 373) and the Francis Crick Institute at 37.6% (157 of 418)—intergovernmental laboratories and independent institutes in which data sharing is deeply embedded in the research workflow.

Private philanthropies and charitable foundations were similarly prominent. Genome Canada reached 35.1% (134 of 382 articles), the Howard Hughes Medical Institute 34.6% (422 of 1,219), the Burroughs Wellcome Fund 32.9% (119 of 362), the Fondation ARC pour la Recherche sur le Cancer 32.0% (87 of 272), and the Canadian Institute for Advanced Research (CIFAR) 31.7% (78 of 246)— all roughly three times the 11.7% funder-linked rate.

National research organizations and shared research facilities also exceeded the funder-linked rate by wide margins, including the European Synchrotron Radiation Facility (33.2%), France’s INRAE (29.9%, 161 of 539 articles), and Germany’s Max Planck Society (28.0%, 444 of 1,583) and Leibniz Association (26.7%).

Sub-agency programmes that OpenAlex surfaces as separate funder entities (for example, particular NSF directorates or DOE program offices) are folded into their parent agency following the ROR organizational hierarchy, so the agency rows in Table 1 report the aggregate rather than the constituent. We therefore do not highlight such programmes individually, even where their rates exceed those of their parent agency: isolating the constituents that OpenAlex happens to surface separately would be inconsistent with a design that aggregates constituents to the parent.

### 3.3 Variation by Journal

Journal-level open data rates showed even greater variation than funder rates, spanning from below 5% to above 90% estimated across the full set of 1,388 journals. As with funders, Table 2 and Figure 2 are restricted to high-volume journals by the thresholds of Section 2.5.2; several of the highest-rate journals discussed below are smaller titles that fall below those cutoffs and appear only in the complete rankings released with the analysis code (Data Availability).

**Figure 2:**
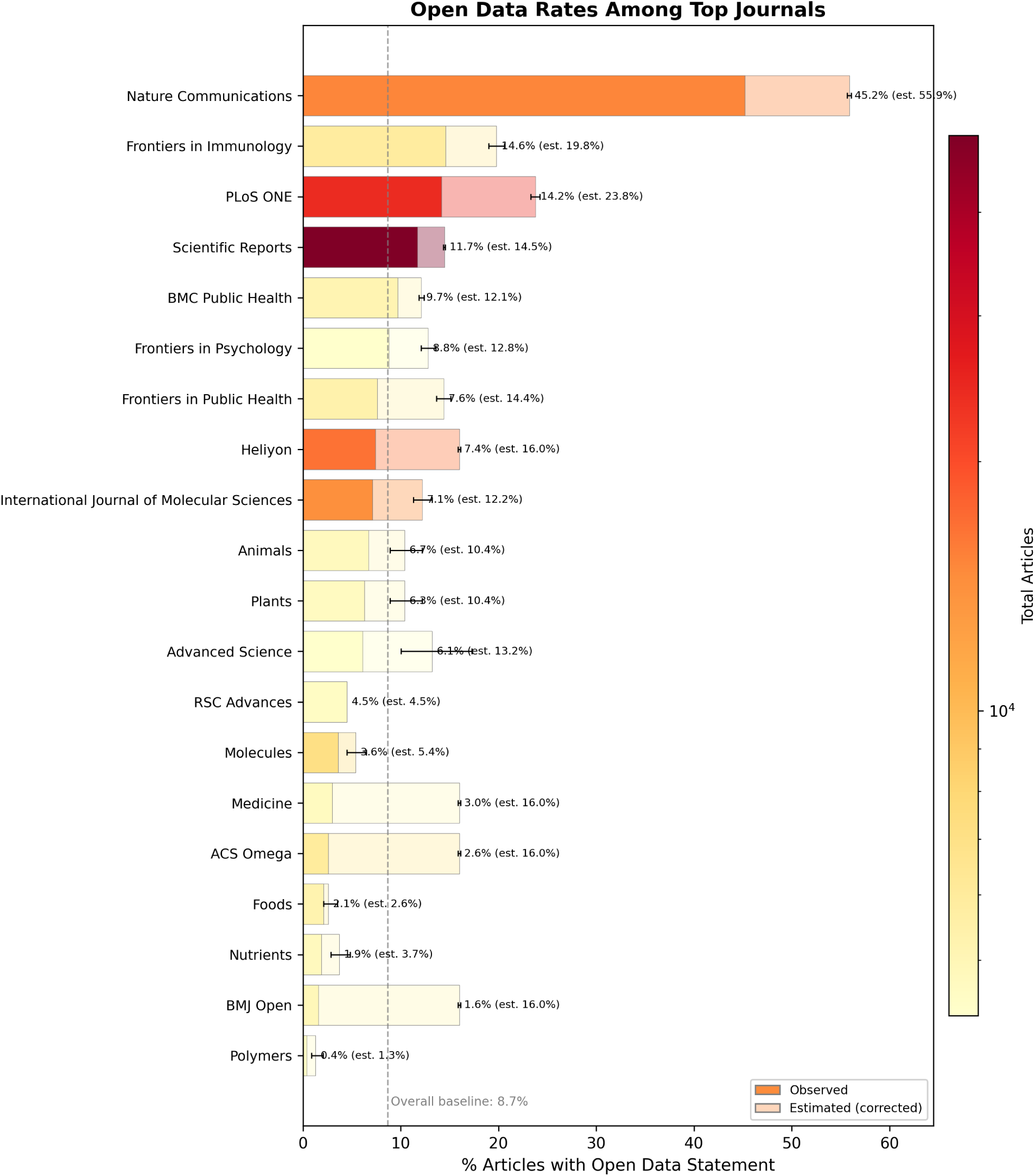
Open data rates among top biomedical journals (2024–2025 research articles). Solid bars show the observed open data rate; lighter background bars show the estimated (corrected) rate after applying journal-level correction factors from head-to-head PDF vs. XML comparison. Error whiskers indicate the 95% imputation interval on the corrected estimate (uncertainty in the XML-only coverage correction, not sampling uncertainty of the observed rate); journals where the correction does nothing carry no whisker. Bar color encodes total article count on a log scale (yellow = fewer, red = more). Dashed line shows the overall baseline. The 20 highest-rate journals among those exceeding the Weibull-derived 2% survival article-count threshold (*≥*4,262 articles) are shown; rankings for the remaining high-volume journals appear in Table 2, and the complete rankings for all 1,388 journals are released with the analysis code.

Journals in genomics and structural biology dominated the top ranks. Nature Structural & Molecular Biology led with an observed rate of 86.1% (130 of 151 articles) and an estimated rate of 91.7% after correction for XML-only coverage. Microbial Genomics (84.8%), Molecular & Cellular Proteomics (81.6%), Cell Genomics (80.2%), and Genome Research (76.0%) all exceeded 75% observed. Nature Genetics achieved the highest estimated rate among all journals at 92.9% (95% imputation interval: 91.5–93.7%), reflecting the substantial uplift from PDF-based detection over its predominantly XML-only coverage. Genome Biology reached an estimated 89.3% (95% imputation interval: 88.1–90.2%).

A notable cluster of psychology and cognitive science journals also demonstrated high open data rates: the Journal of Cognition (72.8%), Attention, Perception, & Psychophysics (62.9% observed, 68.2% estimated), Psychological Research (62.2%), Behavior Research Methods (59.8%), and Communications Psychology (54.1%). These rates co-occur with the growing open data culture in experimental psychology, where community norms and journal requirements emerged in response to the replication crisis in that field.

Among the Nature portfolio, open data rates ranged from 86.1% (Nature Structural & Molecular Biology) to 45.2% (Nature Communications), reflecting the diverse disciplinary scope of these journals and varying levels of field-specific data sharing norms. Nature Immunology (62.8% observed, 79.1% estimated), Nature Cell Biology (62.1% observed, 76.3% estimated), Nature Methods (59.0% observed, 73.7% estimated), and Nature Neuroscience (55.5% observed, 69.8% estimated) all showed substantial upward corrections, indicating that PDF-based detection captured data sharing statements missed by XML extraction in these journals.

High-volume journals provide insight into data sharing at scale. Scientific Data published 2,421 articles in our window at a 65.6% observed open data rate. This figure is best read as a property of our detector rather than of the journal: Scientific Data is a dedicated data journal whose articles are, by editorial design, descriptors of deposited datasets, so its true rate should approach 100%. The observed 65.6% instead sits close to the 61.8% sensitivity for open data detection reported by external validation (*22*), and thus serves as an in-corpus calibration of the detection floor discussed in the Limitations. The rates below should be read with that floor in mind: they are lower bounds on declared sharing, and the gap is largest where a field’s repositories and phrasings are least well covered by the detector’s dictionary. iScience (3,514 articles, 50.8%), Communications Biology (2,503 articles, 47.2% observed, 60.1% estimated), and Nature Communications (15,129 articles, 45.2% observed, 55.9% estimated) demonstrate that moderately high rates are achievable even at very large publication volumes.

### 3.4 Interactive Data Exploration

#### Interactive Dashboard

Readers can explore these results dynamically through our web dashboard at https://www.opensciencemetrics.org, which provides real-time filtering and visualization across multiple dimensions including time periods, countries, and research domains.

## 4 Discussion

### 4.1 Key Findings

Our analysis of 951,949 open access biomedical articles reveals that open data sharing is not uniformly distributed across the research ecosystem. The overall rate of 8.7%—and even the 11.7% rate among funder-linked articles—falls far short of universal compliance with data sharing mandates. Yet the more than tenfold variation we observe, from single-digit percentages to above 90%, shows that high data sharing rates do occur in some funder and journal contexts, even if our cross-sectional design cannot say what drives that variation.

High open data rates at both the funder and journal level co-occur with two factors visible in our data: research domains where data sharing is technically routine (genomics, structural biology, environmental science) and funder or journal contexts with explicit, enforced sharing requirements. Funders with strong institutional cultures of openness (e.g., EMBL, EMBO, the Francis Crick Institute) reach rates three to four times the funder-linked rate. Among journals, the pattern is more pronounced: journals that require data deposit as a condition of publication (e.g., Nature Genetics, Genome Research) reach rates above 75%, while journals with weaker or absent data sharing requirements remain near or below the baseline. Because funders and journals are not randomly assigned to articles, and because policy and discipline are tightly confounded, we cannot attribute these differences to policy text as such.

### 4.2 Comparison to Previous Work

Our results extend and refine previous large-scale assessments of research transparency. Serghiou et al. documented the prevalence of multiple transparency indicators across 2.75 million open access PubMed Central articles published between 1959 and 2020, using XML-based text mining and an early version of the same oddpub detection algorithm (*13*). The appropriate comparison is with our *corpus-wide* rate of 8.7%, not the 11.7% funder-linked rate: Serghiou et al. did not restrict to articles with an identified funder, and comparisons that conflate the two rates will misleadingly inflate any apparent difference. Restricting to research articles, as our corpus does, they estimated a 2020 data sharing rate of 17% (and 15% across all article types that year).

These two figures measure different things, and the gap between them should not be read as a decline in data sharing. Two factors should have moved our estimate *upward* relative to theirs.

Our window falls five years later, over a period in which the trend they documented was rising. And our PDF-based pipeline detects approximately 52% more open data statements than XML-based extraction on the same articles—an improvement attributable to MinerU’s ability to extract text from formatted data availability statements, supplementary sections, and other elements not represented in PMC XML markup—so previous XML-only estimates systematically undercount relative to ours. That the net movement is nonetheless downward points to a third factor operating in the opposite direction and outweighing both: the open data criterion itself became substantially more conservative between the two studies. The version of oddpub available in 2020 credited a supplementary file whose name matched a data-like pattern (for example, “raw data” adjacent to .csv or .xls), a supplementary dataset such as “S1 Table,” or data posted to a code-hosting site. Version 7.2.3 credits none of these, and further separates statements describing *reuse* of existing data from statements describing newly shared data. The criterion thus shifted from detecting essentially any mention of accessible data to detecting only data deposited in a recognized repository with an accession number or persistent link. Much of what a 2020 corpus registered as sharing—files attached to the article itself—is excluded from our rate by construction. Any comparison across the two should be read with that redefinition in mind, and the ordering of effects is itself informative: tightening the definition of open data moved the measured rate more than five additional years of policy and a substantially better text-extraction pipeline moved it the other way.

The correction factors we apply to account for XML-only coverage provide estimated rates that are consistently higher than observed rates, particularly for journals where a large fraction of articles lack PDF extraction. For example, Nature Genetics’ observed rate of 70.5% rises to an estimated 92.9% after correction, reflecting the journal’s strong data sharing culture that is partially masked by incomplete PDF coverage.

The PLOS Open Science Indicators dataset (v10; approximately 139,000 PLOS articles and 28,000 comparator articles with publication dates from 1 January 2018 through 30 March 2025) provides a contemporary external benchmark produced by PLOS in partnership with DataSeer from journal XML (*23*).^1^ The continued use of XML-based extraction in widely cited monitoring datasets underscores the value of the PDF-first approach we adopt for PMC-scale measurement.

Open data detection is also beginning to shift toward large language model (LLM) approaches operating on full text. A March 2026 pilot reported that an LLM applied to a corpus of approximately 6,000 Michael J. Fox Foundation–funded articles detected evidence of data *reuse* in 43% of articles, compared with approximately 2% via traditional identifier-based methods (*24*). That comparison addresses a different task than ours (reuse rather than declared sharing), and the underlying model is closed-source, but it indicates the direction of methodological change. We view keyword- and pattern-based pipelines such as oddpub as the appropriate open-source, auditable, institution-scale baseline against which future LLM-based detectors should be benchmarked.

#### Changes from preliminary results

Earlier versions of this analysis were presented as conference posters, and the funder rates reported here differ from those preliminary numbers because the underlying funder attribution has changed twice. Our first presentation identified funders using a primarily custom named-entity-recognition pipeline over acknowledgement text (*25*); subsequent presentations adopted OpenAlex funder metadata as it stood before the 2026 funding-data expansion (*26*, *27*). The rates reported here use OpenAlex’s expanded 2026 funding data (Methods), which links substantially more articles to each funder—especially at commercial publishers whose funding metadata was previously sparse—than either earlier approach. Because the newly linked articles share data at somewhat lower rates, most per-funder rates are revised modestly downward relative to those preliminary numbers; funders with small initial samples shifted most, as their attributed counts roughly doubled and their rates regressed toward the overall level. Separately, some funders that featured prominently no longer stand out in the main figure. Most notably, the Howard Hughes Medical Institute (HHMI)—a private philanthropy that funds a relatively small number of elite investigators, and one we highlighted in our earliest presentation (*25*)— achieved one of the highest per-article open data rates in the dataset (34.6%, ranked 27th of 1,833 funders). However, HHMI’s 1,219 funder-linked articles in the 2024–2025 window fall well below the Weibull-derived figure threshold of *≥*2,694 articles, and its 40,261 total OpenAlex works fall below the 100,000-work minimum applied to exclude small or niche funders. These thresholds, while necessary for focusing the figure on major world funders, systematically exclude high-performing private foundations with narrow funding portfolios. HHMI and similar funders (e.g., the Simons

Foundation, the Chan Zuckerberg Initiative) remain visible in the full supplementary rankings. This pattern underscores that funder *size* and funder *effectiveness* at promoting open data are distinct dimensions that should be interpreted separately.

#### Validation of funder attribution

Because our funder rankings depend on the completeness of funder attribution, we validated the refreshed OpenAlex attribution against an independent, publisher-sourced benchmark. During the 2026 ICSSI hackathon we cross-referenced our funder identifications against the grant–publication links in Dimensions (*28*), holding each article’s open data outcome fixed and varying only the paper–funder attribution. On the same cohort the two independent sources agree closely: OpenAlex links a funder to 60.2% of articles versus 59.1% for Dimensions, and the resulting funder rankings correlate strongly (Spearman *ρ* = 0.95). This is a marked improvement over the same comparison against our December 2025 attribution (*ρ* = 0.88), reflecting the rapid maturation of OpenAlex’s open funding data over the intervening months. The one funder on which the sources materially disagree is the European Commission, which Dimensions places several points higher on a smaller set of papers; under the more comprehensive attribution it sits mid-pack. Because Dimensions is proprietary and access-restricted, we use it only as an external benchmark, not as a pipeline input: our published rates rest entirely on open, reproducible OpenAlex data that now matches the proprietary source on both coverage and ranking.

### 4.3 Implications for Policy

These results have several implications for research policy, though they should be read as descriptive patterns rather than causal evidence of policy effects. First, funders and journals with explicit, enforced data sharing requirements consistently appear at the top of the rankings, an association compatible with mandates working but also with selection of compliance-friendly researchers into those funders and journals. Second, variation across funders with nominally similar mandates suggests that stringency and enforcement may matter more than the bare existence of a policy — a hypothesis that would need within-funder longitudinal data to test directly. Third, the disciplinary clustering we observe (genomics and psychology journals leading their respective tiers) indicates that community norms co-vary with measured open data rates beyond what any top-down policy text would predict.

Our results also provide a benchmarking framework. Funders can compare their grantees’ data sharing rates against peer organizations, journals can assess their position relative to comparable publications, and institutions can identify areas for targeted investment in data management infrastructure and training. The interactive dashboard at https://www.opensciencemetrics.org is designed to support exactly these comparisons.

These findings are timely given parallel NIH efforts to build researcher-level incentives for data sharing. The Data Sharing Index Challenge, with Phase 2 deliverables due in mid-2026, will produce candidate metrics that score individual investigators on data sharing quality and reuse (*14*, *15*). Whatever metric emerges, validating it at population scale will require independent corpus-derived rates of the type our pipeline produces, suggesting a natural complementarity between top-down incentive design and bottom-up behavioral measurement. The implementation environment is also evolving: NIH released NOT-OD-26-046 in February 2026, introducing a structured (largely yes/no) Data Management and Sharing Plan format effective May 25, 2026 (*29*). While the underlying 2023 DMS Policy is unchanged and this format change affects pre-award plans rather than the published-article content our pipeline measures, it is likely to make pre-award compliance signals more amenable to machine extraction and easier to link to retrospective sharing rates of the kind reported here.

### 4.4 Limitations

Several limitations should be noted. First, our corpus is restricted to open access articles—overwhelmingly those deposited in PubMed Central, together with a small number reached only through a publisher-hosted open access PDF; closed-access articles are absent entirely and may exhibit different data sharing patterns, likely biased toward lower rates. Second, external validation at the University of Edinburgh (*22*) found high precision for detecting data sharing statements, but sensitivity of only 61.8% for open data detection even when a data availability statement was present—and 52% without that restriction—materially below the 0.73 sensitivity reported in the original 2020 validation (*19*). Sensitivity for open code was lower still, at 30%. Our reported rates therefore likely underestimate true declared sharing, even before considering non-standard phrasings or implicit data sharing (e.g., data available “upon request”). Third, detecting a data sharing *statement* is not equivalent to verifying that data are actually accessible and reusable; our rates should be interpreted as upper bounds on genuine data availability. A systematic review with meta-analysis of the medical and health sciences literature found that, between 2016 and 2021, 8% (95% CI 5–11%) of studies declared public data availability while only 2% (1–3%) actually made data available (*30*); an audit of 3,191 papers across four leading medical journals (BMJ, JAMA, NEJM, Lancet) reached similar conclusions about the gap between stated policy and author practice (*31*). This statement-versus-practice gap is roughly fourfold, and it compounds rather than offsets the sensitivity limitation above: our rates understate declared sharing and overstate genuine availability. Fourth, the correction factors assume that the head-to-head subset (articles with both PDF and XML) is representative of XML-only articles within each journal, which may not hold uniformly. This assumption is weakest precisely where it does the most work: 738 of the 1,388 journals analyzed (31% of the journal-level corpus) have too little head-to-head coverage to support a journal-specific factor and instead receive the 16.0% global fallback, so their corrected estimates carry no within-journal evidence at all. For journals whose subject matter makes the corpus average implausible—clinical case-report titles, for example, where observed rates are near zero—the corrected estimate should be treated as an artifact of the fallback rather than as an estimate of that journal’s practice, and the observed rate is the more informative quantity. We therefore report observed rates as the headline throughout, and a reader comparing journals should prefer them. Fifth, our January 2024–June 2025 window captures a snapshot of current practices but does not permit longitudinal analysis of trends over time. Sixth, article-type filtering removes correction and erratum notices but does not independently exclude retracted articles that remain typed as research articles; given the low base rate of retractions, the effect on aggregate rates is negligible. Seventh, and most fundamentally, the analysis is cross-sectional with no counterfactual: funders and journals are not randomly assigned to articles, so any rate differences across them reflect a confounded mixture of policy text, enforcement, discipline, infrastructure, and researcher self-selection. Distinguishing the contribution of any one factor would require either a longitudinal pre/post design around a specific policy change or a quasi-experimental comparison with explicit controls.

### 4.5 Future Directions

Future work will increase PDF extraction coverage for the XML-only portion of the current 951,949-article corpus and extend the temporal window beyond June 2025. Longitudinal analysis will enable assessment of whether specific policy changes (e.g., the 2023 NIH Data Management and Sharing Policy) are associated with measurable increases in data sharing rates. Institution-level and data-repository-level analyses are reserved for a follow-up paper, where the substantial methodological work of name normalization (for institutions) and repository-mention extraction (for data repositories) can be treated in the depth they require. A further open question is the head-to-head performance of contemporary LLM-based detectors against the oddpub regex/keyword pipeline on the binary “is open data” task; no such benchmark has yet been published, and the corpus and head-to-head subset described here are well suited to serve as the evaluation substrate.

### 4.6 Conclusion

This study demonstrates that open data sharing in biomedicine varies more than tenfold across funders and journals. Many major funders exceeded the 11.7% funder-linked rate, with top-ranked organizations reaching observed rates above 40%; leading journals reached estimated rates above 90%, compared to an overall rate of 8.7%. These findings are consistent with policy mandates and community norms being associated with higher data sharing, while also highlighting the considerable gap between current practice and universal compliance. Continued investment in scalable measurement tools, such as the pipeline and dashboard presented here, will be essential for monitoring progress toward a more open and reproducible research enterprise.

## Acknowledgments

This research was supported in part by the Intramural Research Program of the National Institutes of Health (NIH). The contributions of the NIH author(s) were made as part of their official duties as NIH federal employees, are in compliance with agency policy requirements, and are considered Works of the United States Government. However, the findings and conclusions presented in this paper are those of the author(s) and do not necessarily reflect the views of the NIH or the U.S. Department of Health and Human Services (HHS). NIH support was provided under NIMH project ZICMH002960. This project has been funded in whole or in part with federal funds from the National Cancer Institute, National Institutes of Health, under Contract No. 75N91019D00024. The content of this publication does not necessarily reflect the views or policies of the Department of Health and Human Services, nor does mention of trade names, commercial products, or organizations imply endorsement by the U.S. Government.

We thank John Lee, Travis Riddle, David Siedband, Eric Earl, Jason Priem, Heather Piwowar, Kendra Oudyk, Francisco Pereira, Juan Antonio Lossio-Ventura, David Kennedy, and the Tanenbaum Open Science Institute (TOSI) for contributions, comments, and support on earlier versions of this project.

This work utilized the computational resources of the NIH HPC Biowulf cluster (https://hpc.nih.gov).

## Data Availability

Full results—open data rates for all 1,833 funders and 1,388 journals—are available as supplementary CSV files in the manuscript repository at https://github.com/nimh-dsst/osm-preprint-2026, together with the analysis code that generates every table and figure in this paper and the full revision history of the manuscript itself. The interactive dashboard at https://www.opensciencemetrics. org provides dynamic exploration of these results; its source code is available at https://github. com/nimh-dsst/osm-dashboard. Open data detection used oddpub v7.2.3 (*19*), an open-source R package available at https://github.com/quest-bih/oddpub; PDF text extraction used MinerU (*18*). The high-performance computing workflows that apply this detection pipeline across the full corpus are specific to our compute environment and are being prepared for public release.

## Use of Generative AI

The authors used generative AI tools during the preparation of this manuscript, including Anthropic’s Claude (via claude.ai and Claude Code), OpenAI models, and the Cursor agentic code editor. These tools were used as drafting, editing, and coding assistants: initial prose for several sections was generated from the author’s outlines, analysis notes, and pipeline outputs, then reviewed, revised, and edited by the author over the course of manuscript preparation. AI assistants were also used during development of the analysis pipeline and supporting code. The author verified all numerical results against the underlying analysis outputs, confirmed that all cited references exist and support the claims made, and is responsible for the accuracy and integrity of the final text. Generative AI was not used to fabricate, alter, or substitute for the underlying data. The full revision history of this manuscript, including every AI-assisted edit, is available in its public Git repository https://github.com/nimh-dsst/osm-preprint-2026.

## A Supplementary Tables

**Table 1:**
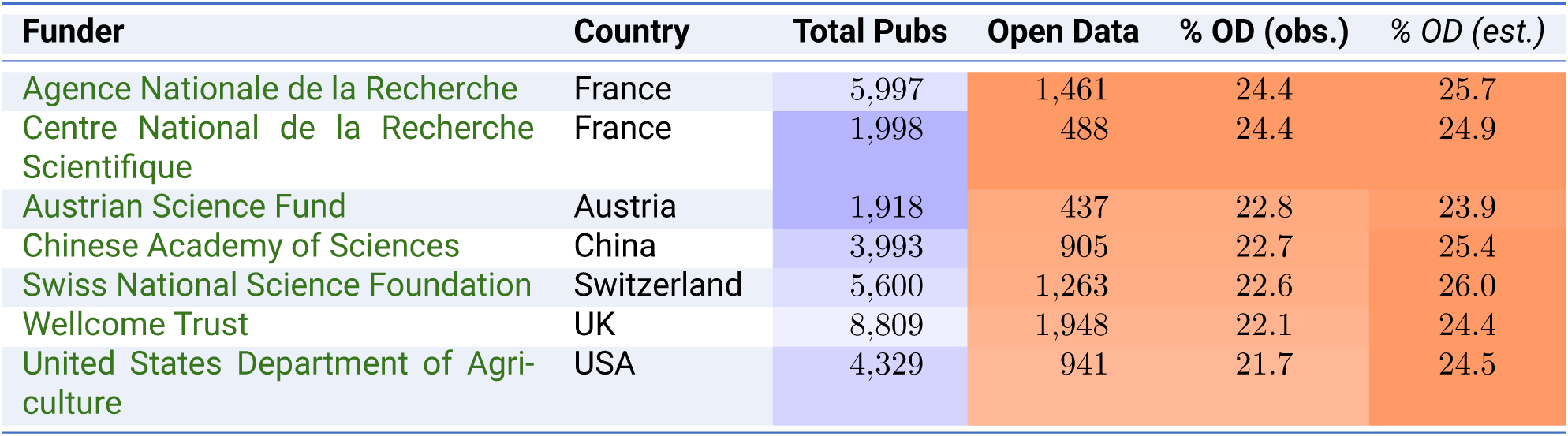
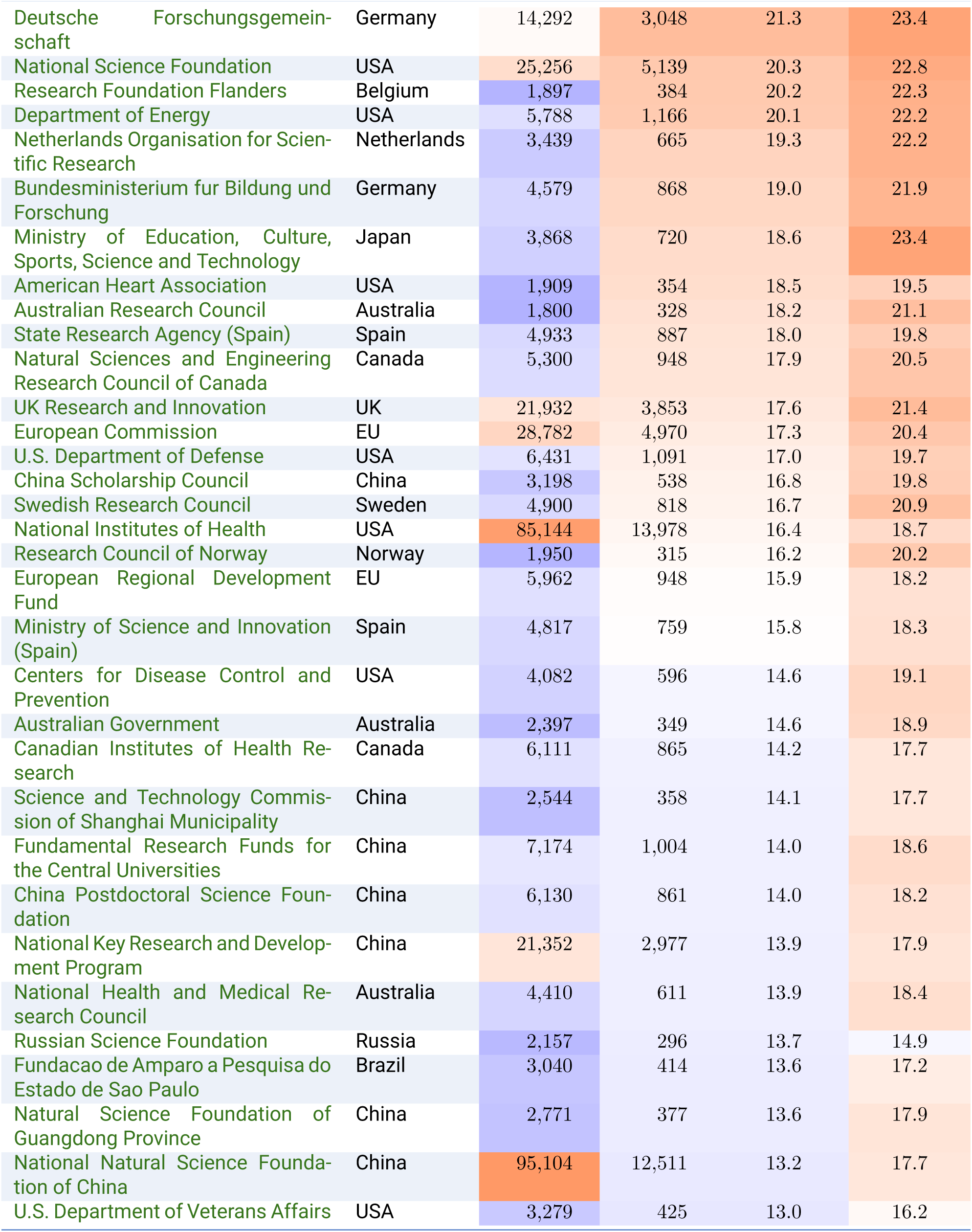
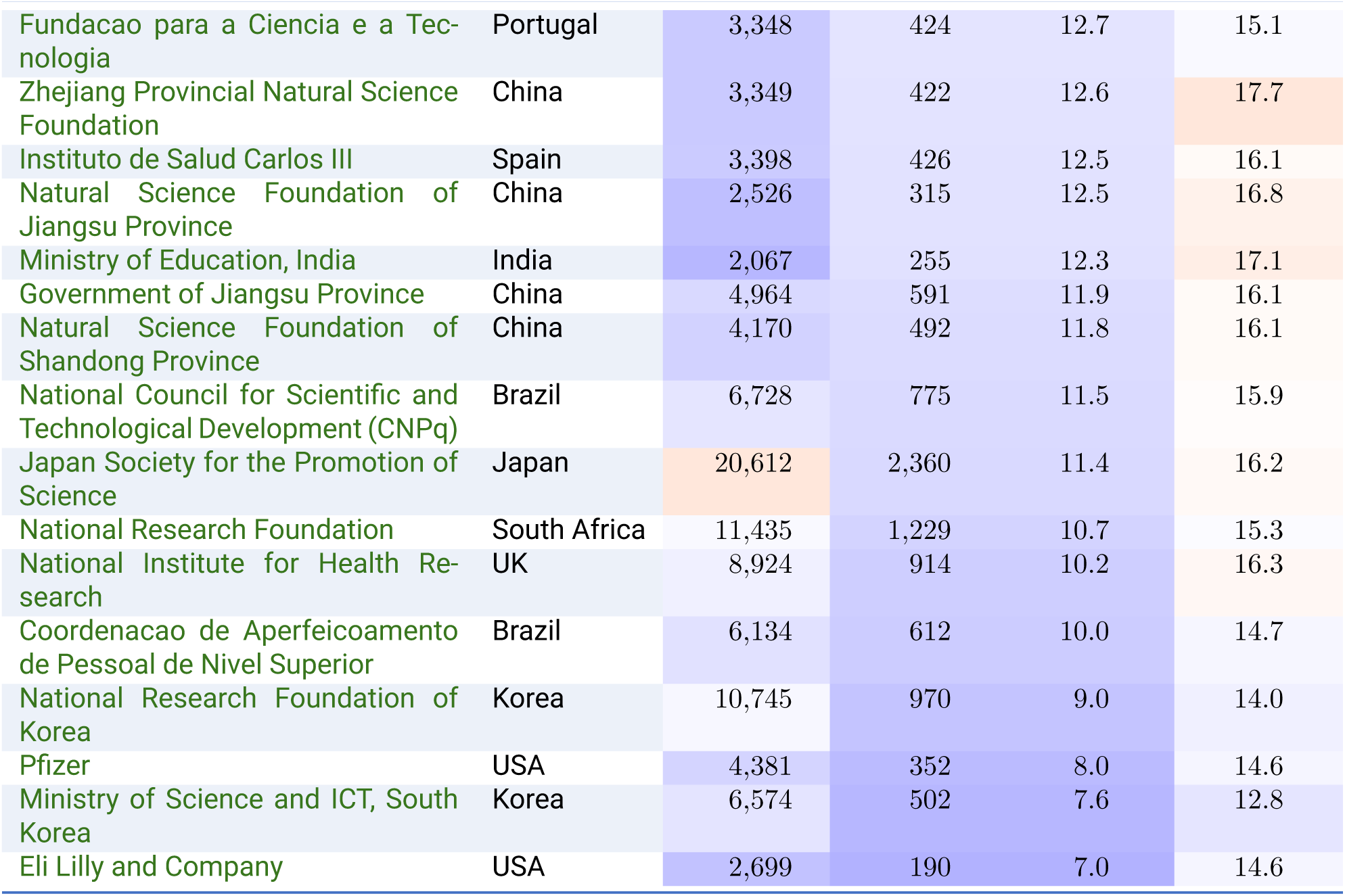
Open data rates among major biomedical research funders. Funders exceeding both the Weibull-derived 5% survival threshold for total funded articles (*≥*1,708 articles with oddpub v7 coverage) and 50,000 aggregated OpenAlex works, ranked by observed open data rate. Parent funders (e.g., NIH, UKRI) aggregate all child institutes with deduplicated article counts; sub-agency programmes (e.g., NSF directorates, the DOE Office of Science) are folded into their parent agency following the ROR organizational hierarchy, so an article crediting both a programme and its agency is counted once. **% OD (obs.)** is the headline rate: the directly measured open data rate across all articles in each funder’s portfolio. *% OD (est.)* is a supplementary modeled estimate that applies journal-level head-to-head PDF vs. XML correction factors to the XML-only portion; it is informative about the magnitude of XML undercount but assumes the head-to-head subset is representative of XML-only articles within each journal. Cell shading: Total Pubs uses a blue-to-red gradient on log scale; both % OD columns share a linear blue-to-red gradient anchored on the observed range. Full rankings for all 1,833 funders are available in the supplementary materials on GitHub.

| Funder | Country | Total Pubs | Open Data | % OD (obs.) | % OD (est.) |
| --- | --- | --- | --- | --- | --- |
| Agence Nationale de la Recherche | France | 5,997 | 1,461 | 24.4 | 25.7 |
| Centre National de la Recherche Scientifique | France | 1,998 | 488 | 24.4 | 24.9 |
| Austrian Science Fund | Austria | 1,918 | 437 | 22.8 | 23.9 |
| Chinese Academy of Sciences | China | 3,993 | 905 | 22.7 | 25.4 |
| Swiss National Science Foundation | Switzerland | 5,600 | 1,263 | 22.6 | 26.0 |
| Wellcome Trust | UK | 8,809 | 1,948 | 22.1 | 24.4 |
| United States Department of Agriculture | USA | 4,329 | 941 | 21.7 | 24.5 |

| **Funder** | **Country** | **Total Pubs** | **Open Data** | **% OD (obs.)** | **% OD (est.)** |
| --- | --- | --- | --- | --- | --- |
| Deutsche Forschungsgemeinschaft | Germany | 14,292 | 3,048 | 21.3 | 23.4 |
| National Science Foundation | USA | 25,256 | 5,139 | 20.3 | 22.8 |
| Research Foundation Flanders | Belgium | 1,897 | 384 | 20.2 | 22.3 |
| Department of Energy | USA | 5,788 | 1,166 | 20.1 | 22.2 |
| Netherlands Organisation for Scientific Research | Netherlands | 3,439 | 665 | 19.3 | 22.2 |
| Bundesministerium für Bildung und Forschung | Germany | 4,579 | 868 | 19.0 | 21.9 |
| Ministry of Education, Culture, Sports, Science and Technology | Japan | 3,868 | 720 | 18.6 | 23.4 |
| American Heart Association | USA | 1,909 | 354 | 18.5 | 19.5 |
| Australian Research Council | Australia | 1,800 | 328 | 18.2 | 21.1 |
| State Research Agency (Spain) | Spain | 4,933 | 887 | 18.0 | 19.8 |
| Natural Sciences and Engineering Research Council of Canada | Canada | 5,300 | 948 | 17.9 | 20.5 |
| UK Research and Innovation | UK | 21,932 | 3,853 | 17.6 | 21.4 |
| European Commission | EU | 28,782 | 4,970 | 17.3 | 20.4 |
| U.S. Department of Defense | USA | 6,431 | 1,091 | 17.0 | 19.7 |
| China Scholarship Council | China | 3,198 | 538 | 16.8 | 19.8 |
| Swedish Research Council | Sweden | 4,900 | 818 | 16.7 | 20.9 |
| National Institutes of Health | USA | 85,144 | 13,978 | 16.4 | 18.7 |
| Research Council of Norway | Norway | 1,950 | 315 | 16.2 | 20.2 |
| European Regional Development Fund | EU | 5,962 | 948 | 15.9 | 18.2 |
| Ministry of Science and Innovation (Spain) | Spain | 4,817 | 759 | 15.8 | 18.3 |
| Centers for Disease Control and Prevention | USA | 4,082 | 596 | 14.6 | 19.1 |
| Australian Government | Australia | 2,397 | 349 | 14.6 | 18.9 |
| Canadian Institutes of Health Research | Canada | 6,111 | 865 | 14.2 | 17.7 |
| Science and Technology Commission of Shanghai Municipality | China | 2,544 | 358 | 14.1 | 17.7 |
| Fundamental Research Funds for the Central Universities | China | 7,174 | 1,004 | 14.0 | 18.6 |
| China Postdoctoral Science Foundation | China | 6,130 | 861 | 14.0 | 18.2 |
| National Key Research and Development Program | China | 21,352 | 2,977 | 13.9 | 17.9 |
| National Health and Medical Research Council | Australia | 4,410 | 611 | 13.9 | 18.4 |
| Russian Science Foundation | Russia | 2,157 | 296 | 13.7 | 14.9 |
| Fundacao de Amparo a Pesquisa do Estado de Sao Paulo | Brazil | 3,040 | 414 | 13.6 | 17.2 |
| Natural Science Foundation of Guangdong Province | China | 2,771 | 377 | 13.6 | 17.9 |
| National Natural Science Foundation of China | China | 95,104 | 12,511 | 13.2 | 17.7 |
| U.S. Department of Veterans Affairs | USA | 3,279 | 425 | 13.0 | 16.2 |
| Fundacao para a Ciencia e a Tecnologia | Portugal | 3,348 | 424 | 12.7 | 15.1 |
| Zhejiang Provincial Natural Science Foundation | China | 3,349 | 422 | 12.6 | 17.7 |
| Instituto de Salud Carlos III | Spain | 3,398 | 426 | 12.5 | 16.1 |
| Natural Science Foundation of Jiangsu Province | China | 2,526 | 315 | 12.5 | 16.8 |
| Ministry of Education, India | India | 2,067 | 255 | 12.3 | 17.1 |
| Government of Jiangsu Province | China | 4,964 | 591 | 11.9 | 16.1 |
| Natural Science Foundation of Shandong Province | China | 4,170 | 492 | 11.8 | 16.1 |
| National Council for Scientific and Technological Development (CNPq) | Brazil | 6,728 | 775 | 11.5 | 15.9 |
| Japan Society for the Promotion of Science | Japan | 20,612 | 2,360 | 11.4 | 16.2 |
| National Research Foundation | South Africa | 11,435 | 1,229 | 10.7 | 15.3 |
| National Institute for Health Research | UK | 8,924 | 914 | 10.2 | 16.3 |
| Coordenacao de Aperfeicoamento de Pessoal de Nivel Superior | Brazil | 6,134 | 612 | 10.0 | 14.7 |
| National Research Foundation of Korea | Korea | 10,745 | 970 | 9.0 | 14.0 |
| Pfizer | USA | 4,381 | 352 | 8.0 | 14.6 |
| Ministry of Science and ICT, South Korea | Korea | 6,574 | 502 | 7.6 | 12.8 |
| Eli Lilly and Company | USA | 2,699 | 190 | 7.0 | 14.6 |

**Table 2:**
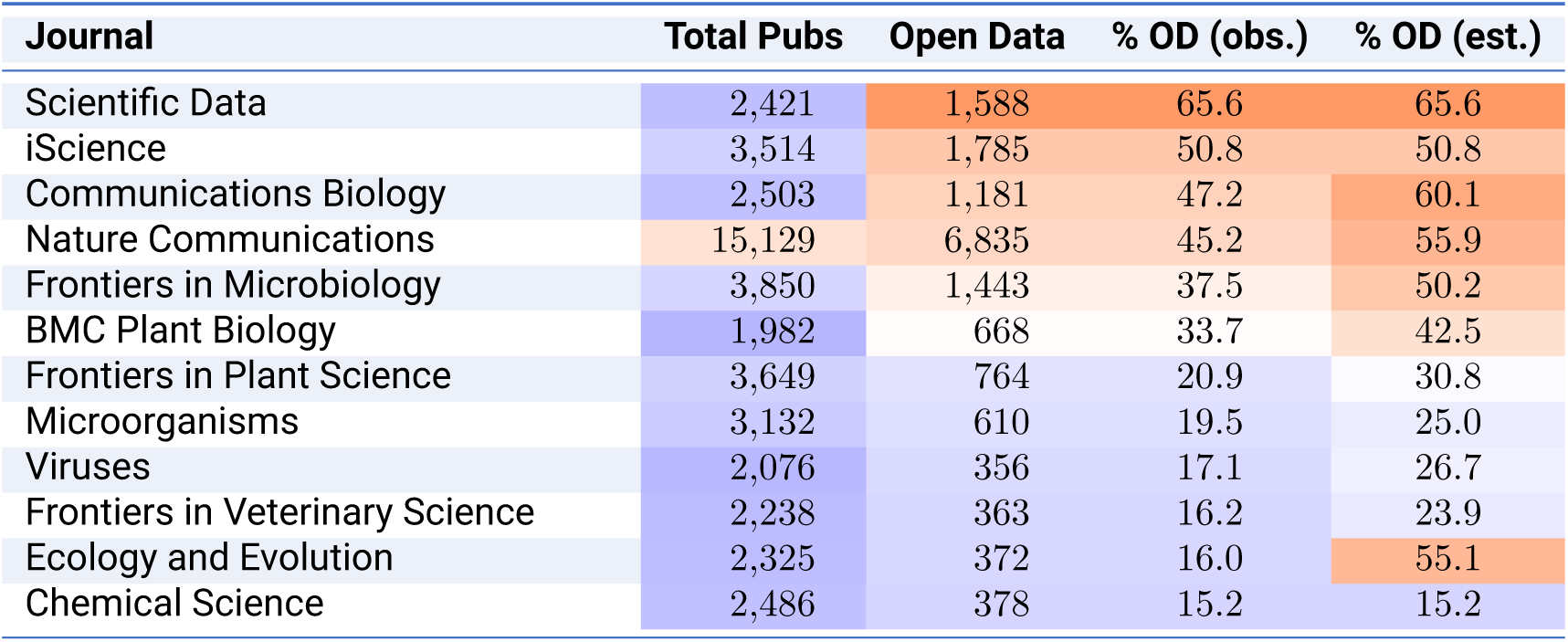
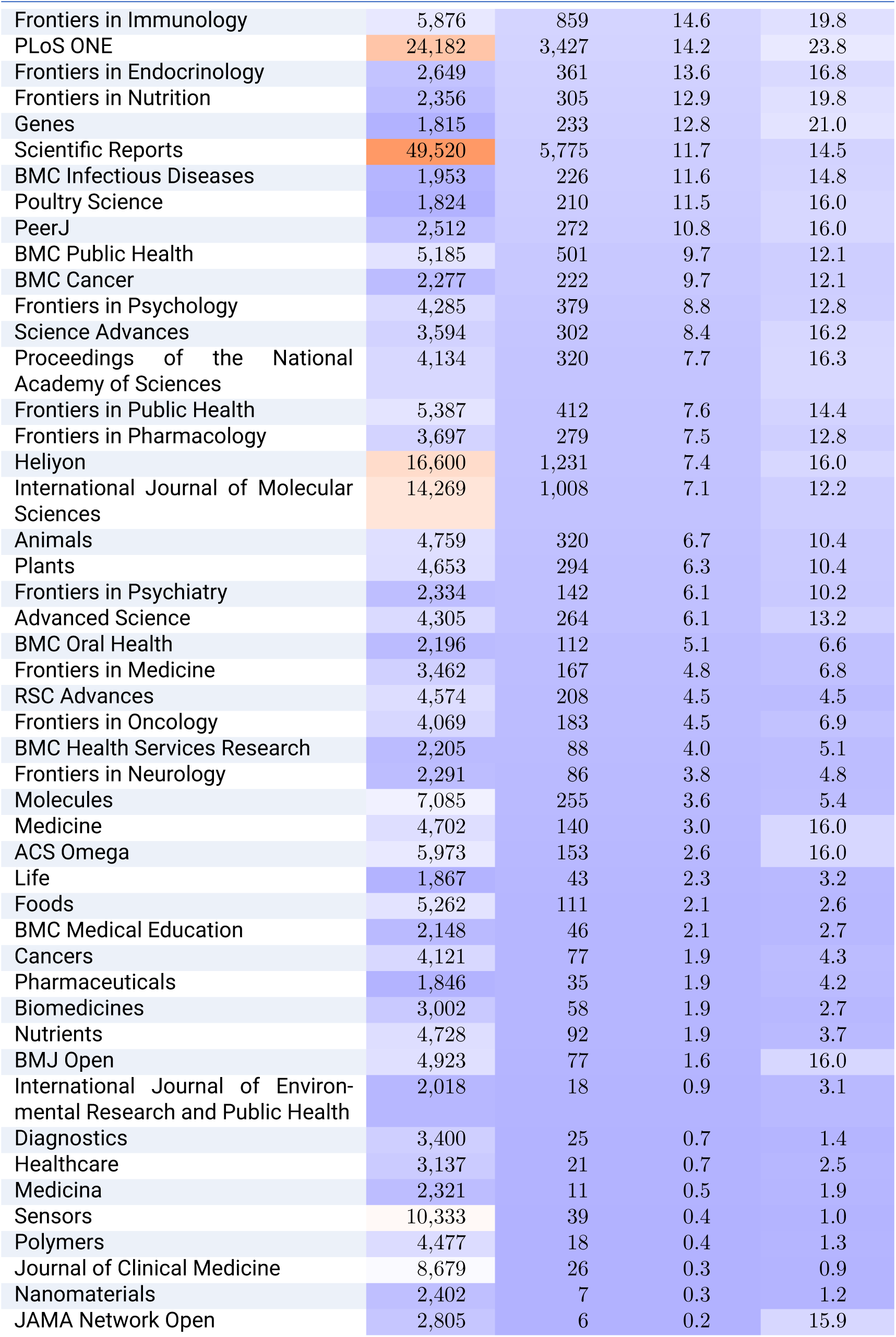
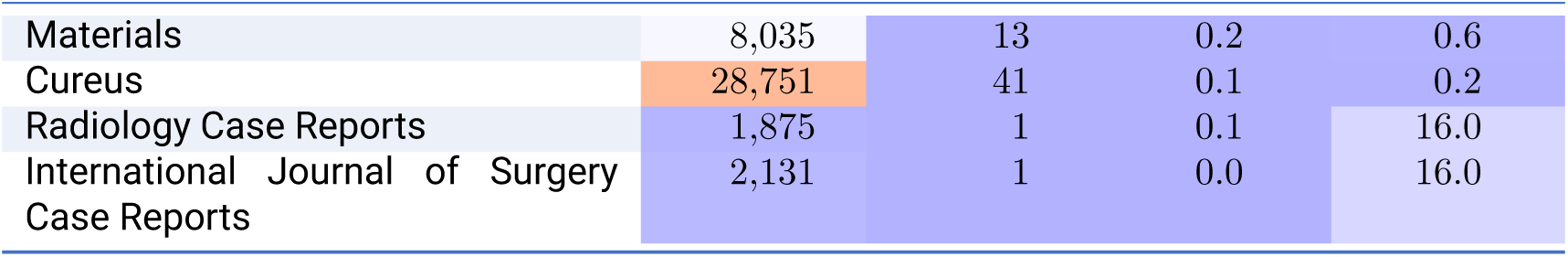
Open data rates among top biomedical journals. Journals exceeding the Weibull-derived 5% survival threshold for total articles (*≥*1,815 articles with oddpub v7 coverage), ranked by observed open data rate. % OD (obs.) shows the directly measured rate; % OD (est.) applies journal-level correction factors from head-to-head PDF vs. XML comparison to estimate the true rate for articles with XML-only coverage. Cell shading: Total Pubs uses a blue-to-red gradient on log scale; % OD columns use a linear blue-to-red gradient. Full rankings for all 1,388 journals are available in the supplementary materials on GitHub.

| Journal | Total Pubs | Open Data | % OD (obs.) | % OD (est.) |
| --- | --- | --- | --- | --- |
| Scientific Data | 2,421 | 1,588 | 65.6 | 65.6 |
| iScience | 3,514 | 1,785 | 50.8 | 50.8 |
| Communications Biology | 2,503 | 1,181 | 47.2 | 60.1 |
| Nature Communications | 15,129 | 6,835 | 45.2 | 55.9 |
| Frontiers in Microbiology | 3,850 | 1,443 | 37.5 | 50.2 |
| BMC Plant Biology | 1,982 | 668 | 33.7 | 42.5 |
| Frontiers in Plant Science | 3,649 | 764 | 20.9 | 30.8 |
| Microorganisms | 3,132 | 610 | 19.5 | 25.0 |
| Viruses | 2,076 | 356 | 17.1 | 26.7 |
| Frontiers in Veterinary Science | 2,238 | 363 | 16.2 | 23.9 |
| Ecology and Evolution | 2,325 | 372 | 16.0 | 55.1 |
| Chemical Science | 2,486 | 378 | 15.2 | 15.2 |
| Frontiers in Immunology | 5,876 | 859 | 14.6 | 19.8 |
| PLoS ONE | 24,182 | 3,427 | 14.2 | 23.8 |
| Frontiers in Endocrinology | 2,649 | 361 | 13.6 | 16.8 |
| Frontiers in Nutrition | 2,356 | 305 | 12.9 | 19.8 |
| Genes | 1,815 | 233 | 12.8 | 21.0 |
| Scientific Reports | 49,520 | 5,775 | 11.7 | 14.5 |
| BMC Infectious Diseases | 1,953 | 226 | 11.6 | 14.8 |
| Poultry Science | 1,824 | 210 | 11.5 | 16.0 |
| PeerJ | 2,512 | 272 | 10.8 | 16.0 |
| BMC Public Health | 5,185 | 501 | 9.7 | 12.1 |
| BMC Cancer | 2,277 | 222 | 9.7 | 12.1 |
| Frontiers in Psychology | 4,285 | 379 | 8.8 | 12.8 |
| Science Advances | 3,594 | 302 | 8.4 | 16.2 |
| Proceedings of the National Academy of Sciences | 4,134 | 320 | 7.7 | 16.3 |
| Frontiers in Public Health | 5,387 | 412 | 7.6 | 14.4 |
| Frontiers in Pharmacology | 3,697 | 279 | 7.5 | 12.8 |
| Heliyon | 16,600 | 1,231 | 7.4 | 16.0 |
| International Journal of Molecular Sciences | 14,269 | 1,008 | 7.1 | 12.2 |
| Animals | 4,759 | 320 | 6.7 | 10.4 |
| Plants | 4,653 | 294 | 6.3 | 10.4 |
| Frontiers in Psychiatry | 2,334 | 142 | 6.1 | 10.2 |
| Advanced Science | 4,305 | 264 | 6.1 | 13.2 |
| BMC Oral Health | 2,196 | 112 | 5.1 | 6.6 |
| Frontiers in Medicine | 3,462 | 167 | 4.8 | 6.8 |
| RSC Advances | 4,574 | 208 | 4.5 | 4.5 |
| Frontiers in Oncology | 4,069 | 183 | 4.5 | 6.9 |
| BMC Health Services Research | 2,205 | 88 | 4.0 | 5.1 |
| Frontiers in Neurology | 2,291 | 86 | 3.8 | 4.8 |
| Molecules | 7,085 | 255 | 3.6 | 5.4 |
| Medicine | 4,702 | 140 | 3.0 | 16.0 |
| ACS Omega | 5,973 | 153 | 2.6 | 16.0 |
| Life | 1,867 | 43 | 2.3 | 3.2 |
| Foods | 5,262 | 111 | 2.1 | 2.6 |
| BMC Medical Education | 2,148 | 46 | 2.1 | 2.7 |
| Cancers | 4,121 | 77 | 1.9 | 4.3 |
| Pharmaceuticals | 1,846 | 35 | 1.9 | 4.2 |
| Biomedicines | 3,002 | 58 | 1.9 | 2.7 |
| Nutrients | 4,728 | 92 | 1.9 | 3.7 |
| BMJ Open | 4,923 | 77 | 1.6 | 16.0 |
| International Journal of Environmental Research and Public Health | 2,018 | 18 | 0.9 | 3.1 |
| Diagnostics | 3,400 | 25 | 0.7 | 1.4 |
| Healthcare | 3,137 | 21 | 0.7 | 2.5 |
| Medicina | 2,321 | 11 | 0.5 | 1.9 |
| Sensors | 10,333 | 39 | 0.4 | 1.0 |
| Polymers | 4,477 | 18 | 0.4 | 1.3 |
| Journal of Clinical Medicine | 8,679 | 26 | 0.3 | 0.9 |
| Nanomaterials | 2,402 | 7 | 0.3 | 1.2 |
| JAMA Network Open | 2,805 | 6 | 0.2 | 15.9 |
| Materials | 8,035 | 13 | 0.2 | 0.6 |
| Cureus | 28,751 | 41 | 0.1 | 0.2 |
| Radiology Case Reports | 1,875 | 1 | 0.1 | 16.0 |
| International Journal of Surgery<br>Case Reports | 2,131 | 1 | 0.0 | 16.0 |

1 The OSI release is cumulative from 2018 through March 2025 and covers PLOS journals plus matched non-PLOS comparators; its Data_Shared and related indicators are not directly comparable to oddpub-based rates on our PMC corpus.

## References

1. M. D. Wilkinson, M. Dumontier, I. J. J. Aalbersberg, G. Appleton, M. Axton, A. Baak, N. Blomberg, J.-W. Boiten, L. B. da Silva Santos, P. E. Bourne, J. Bouwman, A. J. Brookes, T. Clark, M. Crosas, I. Dillo, O. Dumon, S. Edmunds, C. T. Evelo, R. Finkers, A. Gonzalez-Beltran, A. J. G. Gray, P. Groth, C. Goble, J. S. Grethe, J. Heringa, P. A. C. ’t Hoen, R. Hooft, T. Kuhn, R. Kok, J. Kok, S. J. Lusher, M. E. Martone, A. Mons, A. L. Packer, B. Persson, P. Rocca-Serra, M. Roos, R. van Schaik, S.-A. Sansone, E. Schultes, T. Sengstag, T. Slater, G. Strawn, M. A. Swertz, M. Thompson, J. van der Lei, E. van Mulligen, J. Velterop, A. Waagmeester, P. Wittenburg, K. Wolstencroft, J. Zhao, B. Mons, en, Sci Data 3, 160018, ISSN: 2052-4463, DOI 10.1038/sdata.2016.18 (Mar. 2016).

2 . M. Baker, Nature 533, 452–454, ISSN: 0028-0836,1476-4687, DOI 10.1038/533452a (May 2016).

3. G. Colavizza, I. Hrynaszkiewicz, I. Staden, K. Whitaker, B. McGillivray, en, PLoS One 15, e0230416, ISSN: 1932-6203, DOI 10.1371/journal.pone.0230416 (Apr. 2020).

4. National Institutes of Health, Final NIH Policy for Data Management and Sharing, Notice NOT-OD-21-013, 2020.

5 European Commission, Horizon Europe Programme Guide: Open Science Requirements, Brussels, 2021.

6. UK Research and Innovation, Concordat on Open Research Data, Swindon, UK, 2016.

7. Wellcome Trust, Policy on Data, Software and Materials Management and Sharing, London, 2017.

8. Deutsche Forschungsgemeinschaft, Guidelines for Safeguarding Good Research Practice: Code of Conduct, Bonn, 2019.

9. Swiss National Science Foundation, Open Research Data (ORD) Policy and Data Management Plan Requirements, Bern, 2017.

10. Agence Nationale de la Recherche, Plan d’action / Science Ouverte: Data Management Plan Requirement for Funded Projects, Paris, 2019.

11. National Science Foundation, Proposal and Award Policies and Procedures Guide (PAPPG), Alexandria, VA, 2023.

12. UNESCO, UNESCO Recommendation on Open Science, Paris, 2021.

13. S. Serghiou, D. G. Contopoulos-Ioannidis, K. W. Boyack, N. Riedel, J. D. Wallach, J. P. A. Ioannidis, PLoS Biol. 19, e3001107, ISSN: 1544-9173,1545-7885, DOI 10.1371 / journal.pbio.3001107 (Mar. 2021).

14. *NIH Challenge aimed at incentivizing data sharing recognizes Phase 1 winners*, en, https://www.nei.nih.gov/research-and-training/research-news/nih-challenge-aimed-incentivizing-data-sharing-recognizes-phase-1-winners, Accessed: 2026-5-17, Sept. 2025.

15. L. Frank, The S-index Challenge: Develop a metric to quantify data-sharing success, en, https://www.thetransmitter.org/open-neuroscience-and-data-sharing/the-s-index-challenge-develop-a-metric-to-quantify-data-sharing-success/, Accessed: 2026-5-17, Oct. 2024.

16. E. Bobrov, N. Riedel, M. Kip, Quantitative Science Studies 5, 383–407, DOI 10.1162/qss_a_00301 (May 2024).

17 A. Iarkaeva, V. Nachev, E. Bobrov, en, PLoS One 19, e0302787, ISSN: 1932-6203, DOI 10.1371/journal.pone.0302787 (May 2024).

17. B. Wang, C. Xu, X. Zhao, L. Ouyang, F. Wu, Z. Zhao, R. Xu, K. Liu, Y. Qu, F. Shang, L. Bo, C. Zhu, W. Ye, arXiv preprint arXiv:2409.18839, DOI 10.48550/arXiv.2409.18839 (2024).

19. N. Riedel, M. Kip, E. Bobrov, Data Sci. J. 19, 42, ISSN: 1683-1470, DOI 10.5334/dsj-2020-042 (2020).

20. J. Priem, H. Piwowar, R. Orr, *arXiv [cs.DL]*, arXiv: 2205.01833 (cs.DL) (May 2022).

21. E. B. Wilson, J. Am. Stat. Assoc22, 209–212, ISSN: 0162-1459,1537-274X, DOI 10.1080/01621459.1927.10502953 (1927).

22. H. Deeb, S. Creasey, D. L. de Ugarte, G. Strevens, T. Usman, H. Y. Wong, M. A. M. Kutzer, E. Wilson, T. Zieliński, A. J. Millar, en, PLoS One 20, e0328065, ISSN: 1932-6203, DOI 10.1371/journal.pone.0328065 (July 2025).

23. Public Library of Science, *PLOS Open Science Indicators*, 2025, DOI 10.6084/m9.figshare. 21687686.

24. *DataSeer develops AI system to track dataset reuse*, en, https://www.researchinformation.info/news/dataseer-develops-ai-system-to-track-dataset-reuse/, Accessed: 2026-5-17, Mar. 2026.

25. A. Thomas, C. Li, J. Lawrimore, D. Moraczewski, Code and details for Open Science Metrics poster presented at EBRAINS / INCF December 11, 2025, en, Dec. 2025, DOI 10.5281/zenodo.17868245.

26. J. Lawrimore, C. Li, D. Moraczewski, J.-B. Poline, A. Thomas, Who funds open science? Tracking data sharing in publications across major funders (OHBM 2026 poster), 2026, DOI 10.5281/ZENODO.20630153.

27. J. Lawrimore, C. Li, D. Moraczewski, J.-B. Poline, A. Thomas, Who funds open science? Tracking data sharing in publications across major funders (ICSSI 2026 poster), 2026, DOI 10.5281/ZENODO.21042765.

28. Digital Science, *Dimensions Database*, https://www.dimensions.ai, Accessed: 2026-7-5, 2026.

29. National Institutes of Health, Notice of Updates to the Data Management and Sharing Plan Format, Notice NOT-OD-26-046, Feb. 2026.

30. D. G. Hamilton, K. Hong, H. Fraser, A. Rowhani-Farid, F. Fidler, M. J. Page, en, BMJ 382, e075767, ISSN: 0959-8138,1756-1833, DOI 10.1136/bmj-2023-075767 (July 2023).

31. W. Li, X. Liu, Q. Zhang, L. Shi, J.-X. Zhang, X. Zhang, J. Luan, Y. Li, T. Xu, R. Zhang, X. Han, J. Lei, X. Wang, Y. Wang, H. Lan, X. Chen, Y. Wu, Y. Wu, L. Xia, H. Liao, C. Shen, Y. Yu, X. Xu, C. Deng, P. Liu, Z. Feng, C.-J. Huang, Z. Chen, medRxiv, DOI 10.1101/2024.08.29.24312818 (2024).

